# Risk taking to obtain reward: gender differences and associations with emotional and depressive symptoms in a nationally representative cohort of UK adolescents

**DOI:** 10.1101/644450

**Authors:** Gemma Lewis, Ramya Srinivasam, Jonathan Roiser, Sarah-Jayne Blakemore, Eirini Flouri, Glyn Lewis

## Abstract

**Background:** Large population-based studies of neuropsychological factors that characterize or precede depressive symptoms are rare and we know little about gender differences in these associations.

**Methods:** The Millennium Cohort is a representative UK birth cohort of children born between 2000 and 2002. The Cambridge Gambling Task (CGT) was completed at ages 11 (n=12,355) and 14 (n=10,578). Our main exposure was proportion of points gambled, when odds of winning were above chance (risk taking to obtain reward). We also examined how much adolescents adjusted bets as odds of winning increased (risk adjustment). Outcomes were emotional symptoms at age 11 (SDQ) and depressive symptoms at age 14 (sMFQ). We calculated cross-sectional associations, and a longitudinal association from age 11 to 14, using linear regressions before and after adjusting for confounders.

**Results:** Females were less risk taking than males (e.g. females bet 9.22, 95% CI 8.65 to 9.80, percentage points lower at age 11). In univariable models there were cross-sectional associations between risk taking and emotional (age 11) or depressive (age 14) symptoms (e.g. at age 14, a 20-percentage point increase in risk taking was associated with a 0.52 reduction in sMFQ points, 95% CI -.71 to -.33). However, cross-sectional associations were explained by gender differences (at age 14 the association adjusted for gender was: .05, 95% CI -.15 to .25). Longitudinally, there was weak evidence of an association between risk taking and depressive symptoms in females only (a 20-point increase in risk taking at age 11 was associated with a reduction of 0.31 sMFQ points at age 14 (95% CI -.60 to -.02).

**Conclusions:** We found evidence that adolescent females were less likely to take risks than adolescent males. There was no strong evidence of an association between risk taking and emotional and depressive symptoms, after accounting for gender.

**Abbreviations:** **CGT:** Millennium Cohort Study (MCS); Cambridge Gambling Task (CGT); Strengths and Difficulties Questionnaire (SDQ); sMFQ: short Mood and Feelings Questionnaire.

## Introduction

Depression is the leading cause of disease burden worldwide and there are no generally accepted methods of prevention (Vos, 2016). Variation in cognitive functioning is a major source of the disability associated with depressive illness and may be a cause of depression rather than just a consequence (Roiser & Sahakian, 2013). The neuropsychological processes underlying cognitive dysfunction in depression are poorly understood. One theory of depression proposes reduced connectivity between neural circuits for executive control (such as the prefrontal cortex) and neural circuits that respond to rewarding or punishing emotional information (such as the ventral striatum) (Furman, Hamilton, & Gotlib, 2011; Treadway & Pizzagalli, 2014). The precise mechanisms are poorly understood but may involve disrupted translation of reward sensitivity into reward seeking behaviours. If true, alterations in reward circuitry and performance on related tasks would be a risk factor for later depression in longitudinal neuroimaging studies.

This question has proved difficult to study. Neuroimaging studies often use small convenience samples, which may lack statistical power and increase the possibility of an unreliable finding (Button et al., 2013a; LeWinn, Sheridan, Keyes, Hamilton, & McLaughlin, 2017). Small samples can also make findings more difficult to generalise and neuroimaging studies in nationally representative samples are rare. Neuroimaging studies, even when large, have strict exclusion criteria (e.g people with tattoos and claustrophobia). Sample composition probably introduces selection bias and contributes to the poor reproducibility of many neuroimaging findings (LeWinn et al., 2017).

An alternate strategy is to investigate behavioural performance on neuropsychological tasks. These tasks can be included in larger nationally representative cohorts that allow more robust conclusions about cross sectional and longitudinal associations. If performance on a task is associated with depressive symptoms, the neural mechanisms underlying the behavior can then be investigated in neuroimaging studies.

The Cambridge Gambling Task (CGT) assesses decision making and risk taking to obtain rewards. Participants are asked to guess the location of a hidden token, and then gamble a proportion of their points on their decision. If people with depression are less sensitive to rewards (and/or more sensitive to punishments (Eshel & Roiser, 2010)), one would expect a smaller proportion of points to be gambled in the CGT, even when the odds of winning are high. Two cohort studies of children and adolescents have found evidence to support this hypothesis (Forbes, Shaw, & Dahl, 2007; Rawal, Collishaw, Thapar, & Rice, 2013), one of boys aged 10 to 11 (Forbes et al., 2007) and one of children (aged 10 to 18 years) with depressed parents (Rawal et al., 2013). In a smaller case-control study of 15 year olds, there were no differences between depressed adolescents and controls (Kyte, Goodyer, & Sahakian, 2005). In the two cohort studies, placing lower stakes when the odds of winning were high was associated with increased depressive symptoms one year later (Forbes et al., 2007; Rawal et al., 2013).

Depression is twice as common in women than men and this gender difference emerges at around 13 years of age (Joinson, Kounali, & Lewis, 2017). It is possible that cognitive risk factors for depression in adolescence are also more common in females than males. To our knowledge, no neuropsychological studies of adolescent risk taking and reward seeking have investigated gender differences, possibly due to small samples. Three adult studies of the Cambridge Gambling Task have found that men and women do not differ in the overall amount bet, but women exhibit lower risk-adjustment (the extent to which the amount bet is increased as the odds of winning increase (Deakin, Aitken, Robbins, & Sahakian, 2004; van den Bos, Homberg, & de Visser, 2013; van den Bos, Taris, Scheppink, de Haan, & Verster, 2014)).

Existing studies of the CGT and depression have various methodological limitations. They are small (n<200) and may have lacked statistical power (Button et al., 2013b; van den Bos et al., 2013). They also use unrepresentative samples and might have been affected by selection bias. The two cohort studies used short follow-up periods (one year) and one only assessed boys up to the age of 12 (Forbes et al., 2007). Depression is rare in boys until later in adolescence so this study might have lacked statistical power to detect an association. The other study contained very few boys with depression (Rawal et al., 2013).

In this study we used a large birth cohort representative of the UK population to test adolescent gender differences in risk taking to obtain reward, and associations with emotional and depressive symptoms. We compared two sets of cross-sectional analyses, at 11 and 14 years of age. We also investigated the longitudinal association between risk taking to obtain reward at 11 years of age and depressive symptoms at 14 years of age.

## Methods

### Sample

The Millennium Cohort Study (MCS) is an ongoing representative study of 18,552 families and 18,818 children born in England and Wales between September 2000 and August 2001, and in Scotland and Northern Ireland between November 2000 and January 2002 (Joshi & Fitzsimons, 2016). Children were identified through government child benefit records and recruited when they were around 9 to 11 months old. More socially deprived and ethnically diverse areas were oversampled, to increase representation. Six waves of data collection have taken place: when children were around 9 months (MCS1), 3 years (MCS2), 5 years (MCS3), 7 years (MCS4), 11 years (MCS5) and 14 years (MCS6). We used data from ages 11 and 14. At age 11, 13,287 families provided data and at age 14, 11,714 (approximately 61% response rate). We excluded 246 sets of twins and 10 sets of triplets so that data were independent. All families gave written informed consent. Data were obtained from the UK Data Archive. Ethics approval for the MCS was obtained from the UK National Health Service Research Ethics Committee (ref: 11/YH/0203).

### Outcome

#### Adolescent depressive symptoms

The sMFQ is a 13-item self-report measure of DSM-IV depressive symptom severity in the past two weeks and was completed at age 14 (only). Possible scores range from 0 to 26, higher scores indicating more severe symptoms. Reliability was high (Cronbach’s alpha 0.97). We analysed continuous sMFQ scores.

#### Child emotional symptoms

The Strengths and Difficulties Questionnaire (SDQ) consists of five sub-scales, each with five items, assessing emotional symptoms, conduct problems, hyperactivity, peer problems and prosocial behaviour. The SDQ was completed by the parent (mostly mothers) at multiple waves, when children were aged 3 through to 14. We used the emotional symptoms sub-scale as the outcome in our age 11 cross-sectional analyses as there was no depression measure.

### Exposure

The Cambridge Gambling Task was completed on a computer at ages 11 and 14. Participants were shown ten red or blue boxes at the top of the screen. At the bottom of the screen were two more boxes, labelled either red or blue. Participants were told that a yellow token was hidden underneath one of the boxes at the top of the screen. They then had to guess whether the token was hidden under a red or blue box, and indicate their choice by selecting a box at the bottom of the screen, Figure 1. The task was in five stages. Stage one was a ‘decision-only’ practice stage with four trials: on each trial, the task was to indicate whether the token was hidden under a red or blue box. In the first trial the interviewer demonstrated, and in the remaining three trials the participant practiced.

**Figure 1.**
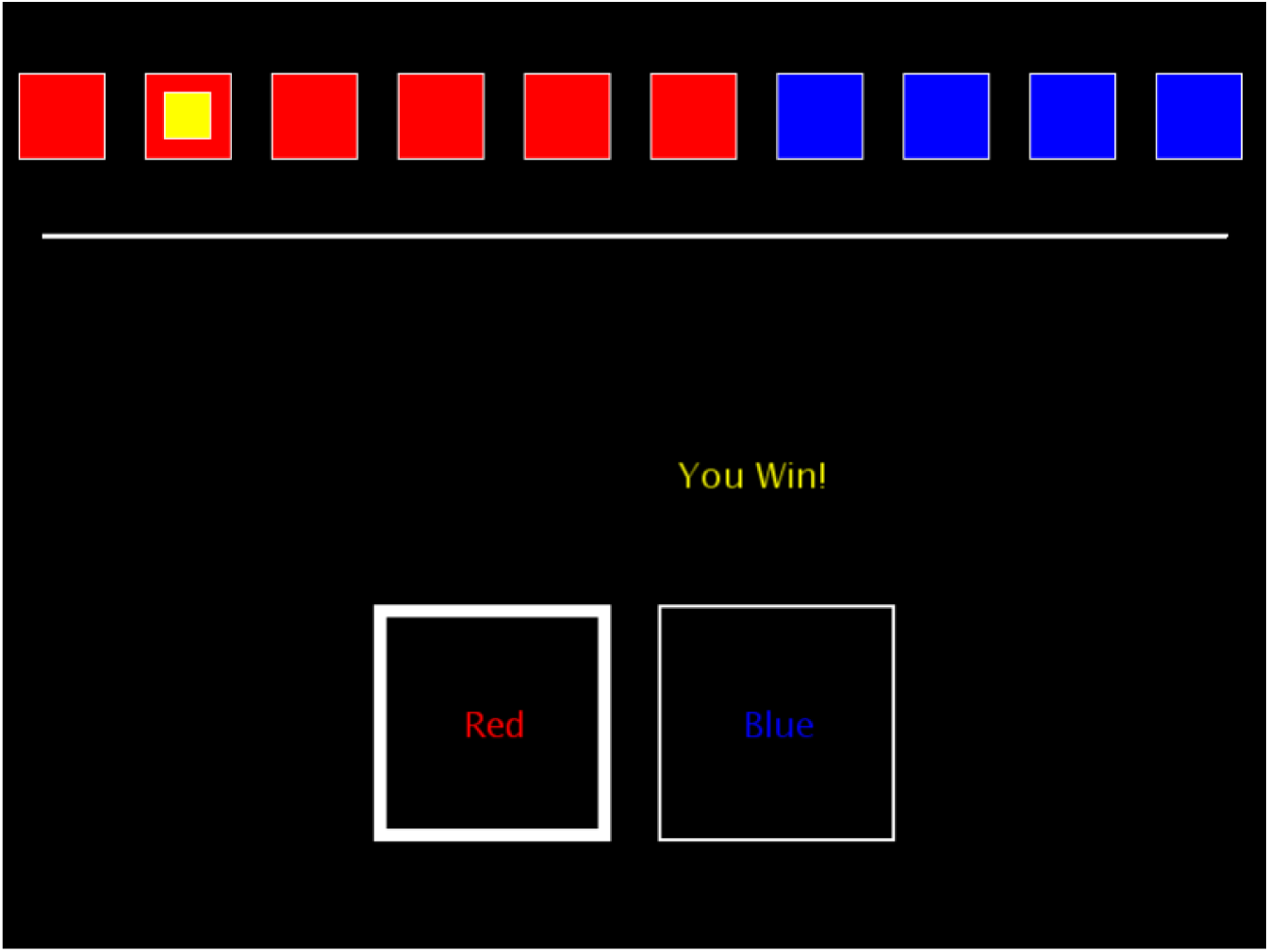
The betting-stage of the Cambridge Gambling Task with a red:blue ratio of 6:4.

After the decision-only stage, the gambling stages (stages two to five) began. At the beginning of each gambling stage, participants were given 100 points and told that their aim was to get as many points as possible before the task ended. After selecting the colour of the box hiding the token, participants were asked to bet a percentage of their points on how confident they were. The percentages were individually flashed onto the screen in five second intervals (5, 25, 50, 75, 95%). The participant selected a percentage and the location of the token was revealed. The value of the bet was either added to (if correct) or subtracted from (if incorrect) the point total (Rawal et al., 2013).

In the first two gambling stages (stages two and three), bets were presented in ascending order (5, 25, 50, 75, 95%). Stage 2 was a practice, with four trials. The interviewer demonstrated the first and the participant practiced the remaining three. The ascending stage then began, with two blocks of nine trials. In the last two gambling stages (stages four and five), bets were presented in descending order (95, 75, 50, 25, 5%; first a practice stage of four trials, then an assessed stage with two blocks of nine trials).

Two outcome variables were derived from the CGT:

1) Risk taking to obtain reward: The mean percentage of the current points total that participants choose to gamble, on trials when they selected the most likely outcome (e.g. trials when they selected a red box, when there were more red than blue boxes). The risk taking variable is therefore the proportion bet (restricted to trials where participants choose the more probable colour), irrespective of the odds of winning (i.e. whether on 6:4, 7:3, 8:2 or 9:1 trials). The percentages that participants can choose from range from 5 to 95% so the risk taking measure ranges from 0.5 to .95. We converted these proportions to percentages (continuous score ranging from 5% to 95%).

2) Risk adjustment: Participants should bet a higher proportion of their points on a certain colour when more of the boxes are that colour. For example, participants should bet more on a red box when the ratio of boxes is 9:1 (red:blue) than when it is 6:4 (red:blue). Risk adjustment measures the extent to which, on trials where a larger proportion of boxes are a certain colour, participants bet a higher proportion of their points. Higher risk adjustment scores represent a higher proportion of points bet as ratio increases. Risk adjustment is calculated using the following formula:

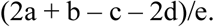

a = mean proportion bet where chosen colour ratio is 9:1

b = mean proportion bet where chosen colour ratio is 8:2

c = mean proportion bet where chosen colour ratio is 7:3

d = mean proportion bet where chosen colour ratio is 6:4

e = mean proportion risked over all trials

#### Potential confounders

We identified variables that might be alternative explanations of observed associations observed between CGT variables and emotional/depressive symptoms. These were variables likely to be associated with both exposure and outcome that were not on the causal pathway. We included family income, maternal education, child age at the time of the exposure, ethnic background, stage of pubertal development, child cognitive ability (as a proxy for intelligence quotient), parent depressive symptoms, and, in longitudinal analyses, children’s baseline emotional symptoms and behavioural problems. Family income was total income adjusted for household size and composition, in quintiles. Maternal education ranged from 1 (none or compulsory education) to 6 (postgraduate), classified into compulsory and non-compulsory. Ethnic background was classified as white or ethnic minority due to small numbers. Pubertal development was stage of breast development in girls and facial hair development in boys (parent report at age 11 and adolescent report at age 14). At age 11, children completed the Verbal Similarities test from the British Abilities Scale assessing verbal reasoning and knowledge. At age 14 they completed a task that measured their understanding of word meanings (10). Parent depressive symptoms were measured using the Kessler 6-item psychological distress scale (K-6) (11). In longitudinal analyses we adjusted for the parent reported SDQ total difficulties score, the sum of the scores on emotional, conduct, hyperactivity and peer problem scales.

### Statistical analyses

All analyses were conducted in Stata version 14, weighted to account for the MCS sampling design and representativeness (using a population weight). In longitudinal analyses we used a non-response weight in addition to the population weight. All analyses are presented for the sample overall, and stratified by gender.

### Gender differences in risk taking and risk adjustment

Gender differences in risk taking and risk adjustment were examined at ages 11 and 14 using independent t-tests. We also calculated the correlation between risk taking at ages 11 and 14, and did the same for risk adjustment.

### Cross-sectional associations at 11 and 14 years of age

First we tested a univariable model with risk taking (continuous exposure) and emotional symptoms (continuous outcome) at 11 years of age. Next we adjusted this model for potential confounders assessed at the time of the exposure or as close as possible to it. Next we adjusted for gender. We also calculated an interaction between gender and risk taking, to test whether the association varied by gender. The same analyses were conducted at age 14, with depressive symptoms as the outcome. We conducted a sensitivity analysis of the age 14 cross-sectional association with SDQ emotional symptoms the outcome. We repeated all analyses with risk adjustment the exposure

### Longitudinal associations from 11 to 14 years of age

First we tested a univariable model with risk taking (continuous exposure) at age 11 and depressive symptoms (continuous outcome) at age 14. Next we adjusted for potential confounders, and then for gender. We also calculated an interaction between gender and risk taking. These analyses were repeated with the age 11 risk adjustment variable the exposure. We conducted a sensitivity analysis with SDQ emotional symptoms the outcome.

### Missing data

Our primary analyses used a complete case sample, with a non-response weight for potential attrition biases in longitudinal analyses. As a sensitivity analysis we report results from an imputed sample. We assumed missingness was dependent on observed data (missing at random), and imputed 50 datasets using multiple imputation by chained equations (MICE). To predict missing data we used all variables selected for analysis models, and a number of auxiliary variables including SDQ measures at all time-points. In cross-sectional analyses, we imputed up to the sample with complete data on the CGT at each time-point (n= 12,355 at age 11 and n= 10,578 at age 14). In longitudinal analyses we used the sample with complete data on the CGT at 11 years of age (n=12,355) and imputed all other variables (confounders and outcome).

## Results

### Descriptive statistics

At age 11, the Cambridge Gambling Task was completed by 12,355 adolescents (mean age 10.67 years, SD .48), 50% female. Data on the outcome (emotional symptoms) were available for 11,931 (97%) of these adolescents. When all confounders were included, we had complete data for 10,396 adolescents (84% of those who completed the CGT). At age 14, the Cambridge Gambling Task was completed by 10,578 adolescents (mean age 13.77 years, SD .45), 50% female. Data on the outcome (depressive symptoms) were available for 10,246 (97%) of these adolescents. When all confounders were included, we had complete data on 8,628 adolescents (82% of those who completed the CGT). Of those who completed the Cambridge Gambling Task at age 11, 9873 (89%) provided MFQ data at age 14. When all confounders were included, we had complete data on a longitudinal sample of 8,418 adolescents (68% of those who completed the CGT at 11 years of age).

Baseline (age 11) characteristics of the sample with complete data are presented in Table 1, according to risk taking (proportion of points bet on trials when the most likely outcome was chosen), split at the median. Children who were less risk taking at age 11 were more likely to be female and more likely to have parents with compulsory education levels only. They were less likely to be in the lowest family income quintile and less likely to be an ethnic minority. Less risk taking children also had lower parental depressive symptoms, lower SDQ total scores at age 11, and a higher mean IQ. At age 14 there was a similar pattern, but no differences according to parental depressive symptoms (Table 2).

**Table 1.**
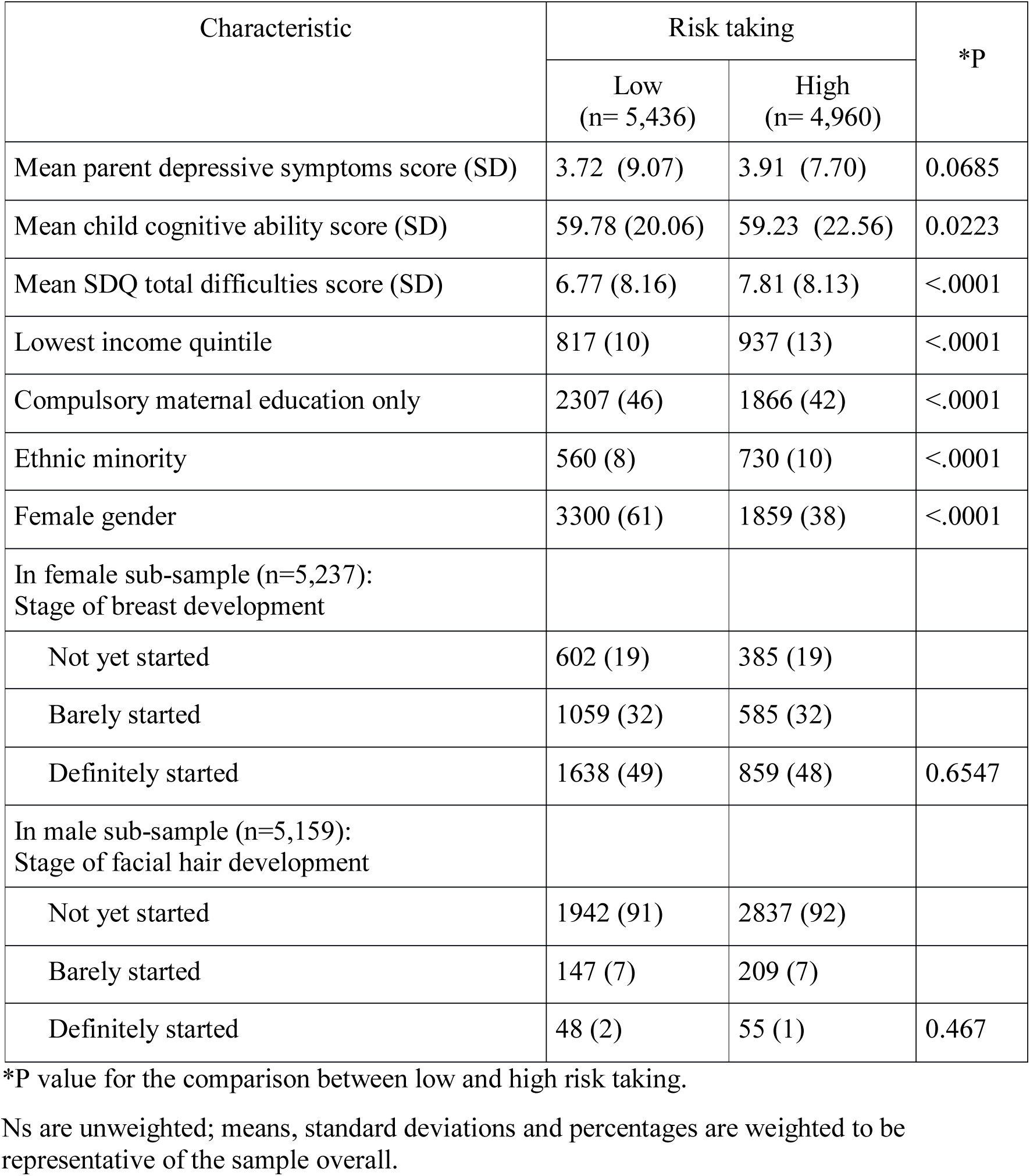
Characteristics of the sample with complete data at 11 years of age, according to reward seeking split at the median (n=10,396). Data are n (%) unless otherwise stated.

**Table 2.**
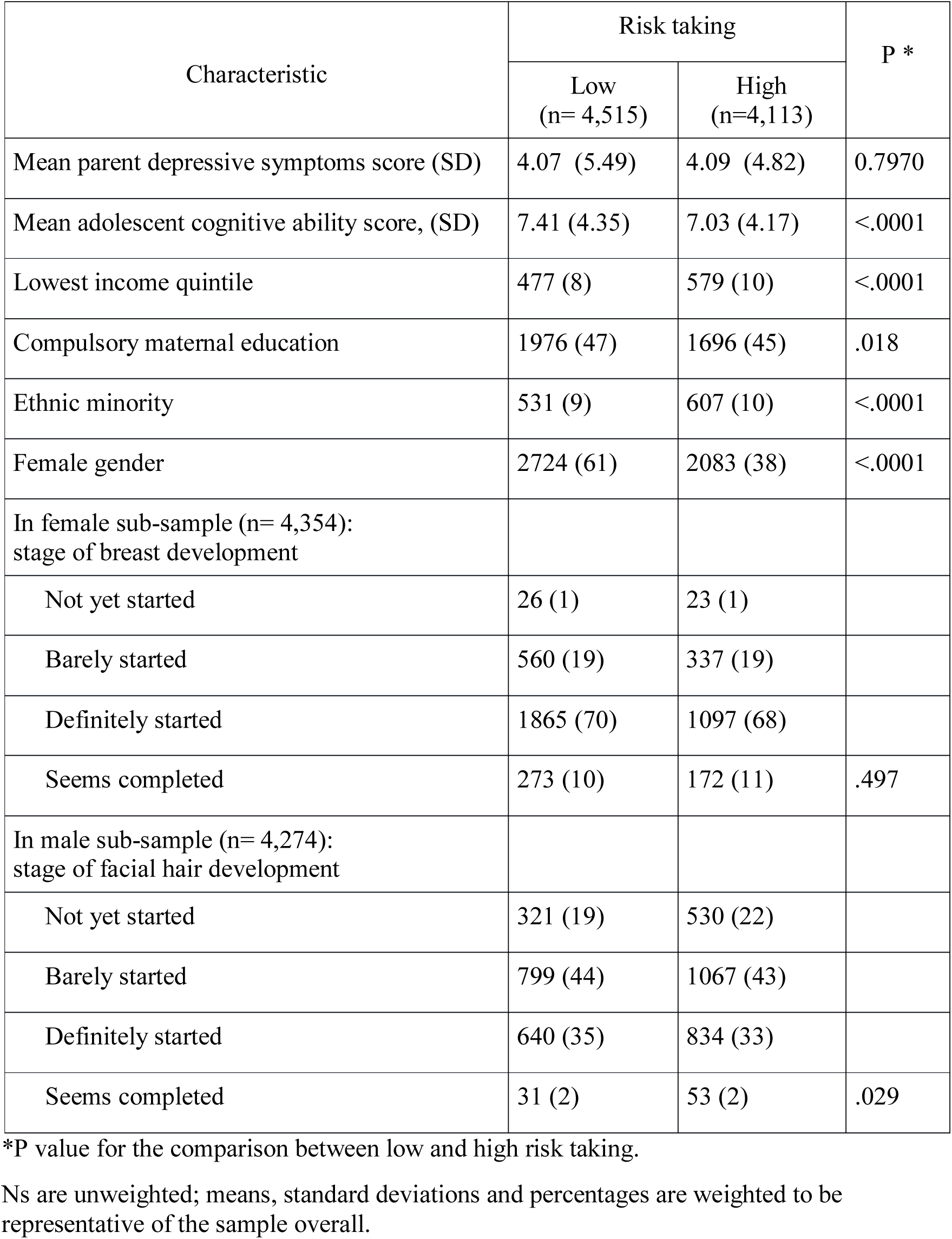
Characteristics of the sample with complete data at 14 years of age, according to risk taking split at the median (n= 8628). Data are n (%) unless otherwise stated.

The mean SDQ emotional symptoms score at 11 years of age was 1.82 (SD) in the sample overall, 1.90 (SD) in females and 1.73 (SD) in males. The mean sMFQ depressive symptoms score at 14 years of age was 5.62 (SD 8.21) in the sample overall, 7.20 (SD 9.52) in females and 4.01 (SD 5.51) in males.

### Gender differences in risk taking and risk adjustment

In the sample overall, the mean risk taking score at 11 years of age was 52.60 (SD 16.92). So, adolescents bet, on average, 53% of their points on trials when they chose the most likely outcome. By 14 years of age, this had decreased slightly to 51.35% (SD 14.67). The correlation between risk taking at ages 11 and 14 in the overall sample was r=.3326 (p<.0001).

At each time-point there was a large gender difference in risk taking, with boys more risk taking than girls (Tables 1 and 2). At 11 years of age, the mean risk taking score for boys was 57. 6% (SD 15.71) and for girls 48.3% (SD 16.75); mean difference 9.22 percentage points (95% CI 8.65 to 9.80, P<.0001). At 14 years of age the mean risk taking score had decreased slightly in boys (55.59%, SD 14.33%) and remained similar in girls (48.10%, SD 14.36%); mean difference 7.49 percentage points (95% CI 6.95 to 8.04, P<.0001). The correlation between risk taking at the two ages was similar in males (r=.2812, p<.0001) and females (r=.2754, p<.0001).

In the sample overall, the mean risk adjustment score at 11 years of age was 0.67 (SD 1.03). By 14 years of age, this had increased to 1.04 (SD 0.98). The correlation between risk adjustment at ages 11 and 14 in the overall sample was r=0.2618 (p<.0001).

At 11 years of age there was no evidence of a gender difference in risk adjustment. The mean risk adjustment score for boys was 0.68 (SD 1.03) and for girls 0.66 (SD 1.04); mean difference 0.03 (95% CI −0.01 to −0.07, P=0.20). At 14 years of age, the mean risk adjustment score had increased in girls and boys but to a larger extent in boys (mean, boys: 1.15, SD 0.98; mean, girls: 0.96, SD 0.97); mean difference 0.20 (95% CI 0.15 to 0.23, P<.0001). The correlation between risk adjustment across time-points was similar in males (r=.2705, p<.0001) and females (r=.2530, p<.0001).

### Cross-sectional associations

Mean SDQ and sMFQ scores according to risk taking and risk adjustment are presented in Supplementary Tables 1 to 3.

Cross-sectional associations between risk taking and emotional symptoms at 11 years of age are presented in Table 3. In a univariable model, we found no evidence of an association (- .03, 95% CI -.08 to .03, p=.314). After adjustments, there was stronger evidence of a negative association (-.07, 95% CI -.12 to -.01, p=.015), indicating negative confounding (Mehio-Sibai, Feinleib, Sibai, & Armenian, 2005). The magnitude of this association reduced after accounting for gender (-.04, 95% CI -.09 to .01, p=.122). When associations were examined separately in males and females (i.e. the potentially confounding role of gender is removed), there was no evidence of an association between risk taking and emotional symptoms, indicating confounding by gender. There was no evidence of an interaction between risk taking and gender (p=0.992).

**Table 3.**
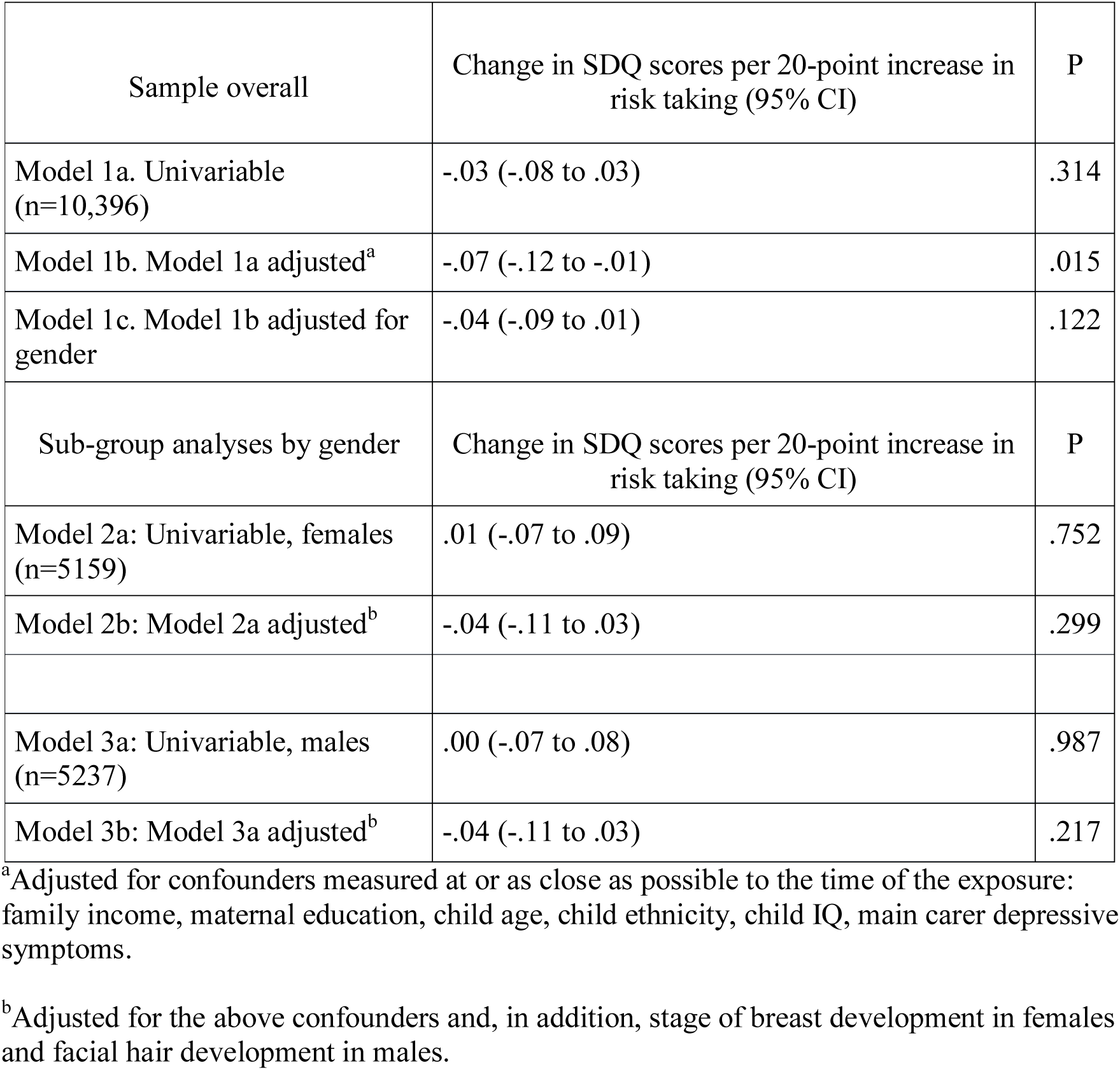
Cross-sectional associations between risk taking (continuous exposure) and emotional symptoms at age 11 (continuous outcome), complete case sample (n=10,396).

We tested for a non-linear association between risk taking and emotional symptoms with a quadratic term in the fully adjusted model. There was no evidence of deviation from linearity (p value for quadratic term .180).

When analyses were repeated with risk adjustment as the exposure, there was no evidence of an association with emotional symptoms after adjustment for confounders (Supplementary Table 9).

Cross-sectional associations between risk taking and depressive symptoms at 14 years of age are presented in Table 4. In the univariable model we found strong evidence of an association. For each 20-point increase in risk taking, depressive symptoms decreased by .52 of a MFQ point (95% CI -.71 to -.33, p<.0001). This was hardly altered after adjusting for confounders (-.54, 95% CI -.73 to -.35, p<.0001) but substantially reduced after accounting for gender (.05, 95% CI -.15 to .25, p=.637). In associations stratified by gender there was no evidence of an association. There was no evidence of an interaction between risk taking and gender (p=0.968).

**Table 4.**
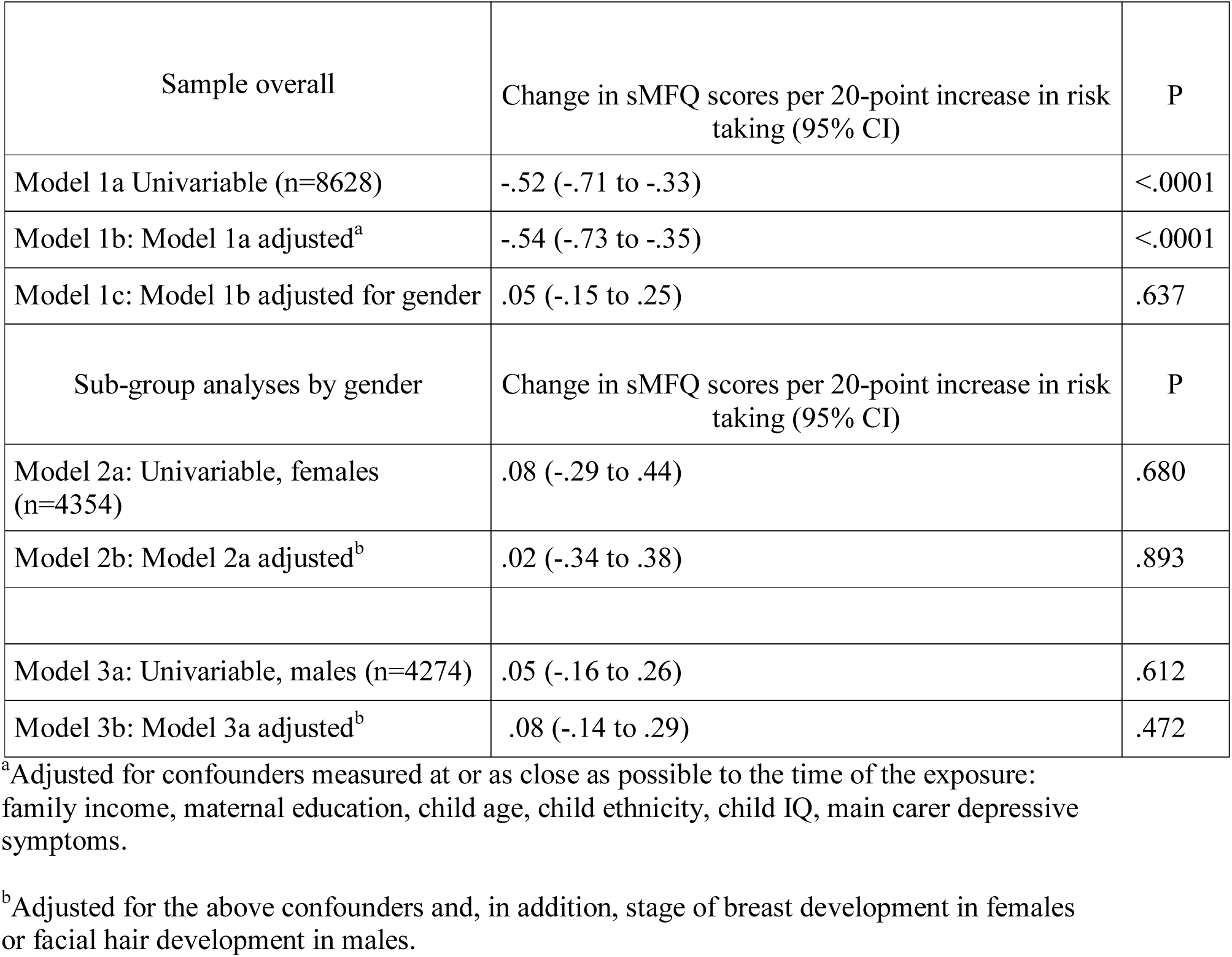
Cross-sectional associations between risk taking (continuous exposure) and depressive symptoms at age 14 (continuous outcome), complete case sample (n=8628).

When analyses were repeated with risk adjustment as the exposure, there was no evidence of an association with depressive symptoms after adjustment for confounders (Supplementary Table 10).

### Longitudinal association

Longitudinal associations are presented in Table 5. In the univariable model we found strong evidence that, for every 20-point increase in risk taking at age 11, depressive symptoms decreased by .55 of a MFQ point at age 14 (95% CI -.76 to -.34, p<.0001). After adjusting for confounders this association was very similar (-.65, 95% CI -.85 to -.44, p<.0001) but it reduced substantially after we further adjusted for gender (-.13, 95% CI -.33 to .07, p=.196). In analyses stratified by gender there was weak evidence of an association in females (-.31, 95% CI -.60 to -.02, p=.037). There was no evidence of an association in males (.11, 95% CI -.11 to .34, p=.328). There was evidence of an interaction between risk taking and gender (p=.015).

**Table 5.** Longitudinal association between risk taking (continuous exposure variable) at age 11 and depressive symptoms (continuous outcome) at age 14, complete case sample with weights for the population and attrition (n=8418).

| Sample overall | Change in sMFQ scores per 20-point increase in risk taking (95% CI) | P |
| --- | --- | --- |
| Model 1a: Univariable (n=8418) | -.55 (-.76 to -.34) | <.0001 |
| Model 1b: Model 1a adjusted<sup>a</sup> | -.65 (-.85 to -.44) | <.0001 |
| Model 1c: Model 1b adjusted for gender | -.13 (-.33 to .07) | .196 |
| Sub-group analyses by gender | Change in sMFQ scores per 20-point increase in risk taking (95% CI) | P |
| Model 2a: Univariable, females (n=4257) | -.27 (-.58 to .04) | .083 |
| Model 2b: Model 2a adjusted<sup>b</sup> | -.31 (-.60 to -.02) | .037 |
| Model 3a: Univariable males (n=4161) | .19 (-.05 to .43) | .121 |
| Model 3b: Model 3a adjusted<sup>b</sup> | .11 (-.11 to .34) | .328 |

<sup>a</sup>Adjusted for confounders measured at or as close as possible to the time of the exposure: family income, maternal education, main carer depressive symptoms, child age, child ethnicity, child IQ and SDQ total difficulties score.
<sup>c</sup>Adjusted for the above confounders and, in addition, stage of breast development in females or facial hair development in males.

With risk adjustment the exposure, there was no evidence of an association with depressive symptoms after adjustment for confounders, Supplementary Table 11.

### Sensitivity analyses

#### Imputed sample

Imputed results for risk adjustment are not presented because this variable showed less evidence of an association with mood in the complete case analyses (but are available on request). A similar pattern of associations for risk taking was observed cross-sectionally and longitudinally in imputed and complete case samples, Supplementary Tables 7 to 9.

#### SDQ at age 14 as the outcome

There was very weak evidence of a cross-sectional association between risk taking and emotional symptoms at 14 years of age after all adjustments including gender (-.06, 95% CI -.12 to .01, p=.091), Supplementary Table 10. There was some evidence of an association in females (-.12, 95% CI -.23 to -.02, p=.022) and very weak evidence of an interaction with gender (.080). Longitudinal associations were similar to those with the MFQ, but there was no evidence of any association in females and no evidence of an interaction with gender, Supplementary Table 11.

## Discussion

We found evidence of a large gender difference in adolescent risk taking to obtain reward, assessed with the Cambridge gambling task. Girls were less risk taking than boys at both 11 and 14 years of age. Reduced risk taking to obtain reward was associated with more emotional symptoms when children were aged 11, and more depressive and emotional symptoms at age 14. However, these associations disappeared after accounting for gender. When cross-sectional associations were run separately in males and females (removing gender as a confounder), there was no evidence of any association between risk taking to obtain reward and emotional/depressive symptoms.

There was strong evidence of a longitudinal association between reduced risk taking at age 11 and depressive symptoms at age 14, but this also disappeared after adjusting for gender. When longitudinal associations were explored separately by gender there was weak evidence that girls (but not boys) who were less risk taking at age 11 had more depressive symptoms at age 14. Statistical evidence for this association was weak and sub-group analyses and interactions can be unreliable, especially when there is no main effect (Rothman, Greenland, & Lash, 2013). In the larger multiply imputed sample, evidence for an interaction and an association in females was weaker, reducing our confidence in this finding.

### Strengths and limitations

This is, to our knowledge, the largest study of risk taking and reward seeking and adolescent depressive symptoms. Our sample was representative of young people in the UK and the cohort design less prone to biases than case-control designs. The three-year follow-up was a strength and included the period during mid-adolescence when depression incidence increases, particularly in females. Our exposures were measured when depression is relatively uncommon, improving the possibility of causal inferences.

Our study has several limitations. Prior longitudinal studies have found that associations were stronger when the likelihood of obtaining a reward was higher (Forbes et al., 2007; Rawal et al., 2013). We could not restrict our analyses to trials with more favourable odds because raw data were unavailable. However, this is measured by risk adjustment, the extent to which participants bet a higher proportion of their points as the probability of winning increases. We found no differences in risk adjustment according to emotional or depressive symptoms. It is therefore unlikely that the probability of winning affected our findings.

In the longitudinal analyses, we adjusted for children’s emotional symptoms and behavioural problems rather than depressive symptoms. The Strengths and Difficulties Questionnaire is a broader concept than depression and is parent-rather than child-reported. Depression is relatively uncommon pre-puberty, so adjusting for a wider range of symptoms at this age could be a better way of accounting for differences in future depression risk, though residual confounding by childhood depression is still possible. Missing data are a limitation of all cohort studies. We used two methods to reduce the possibility of attrition and non-response bias and our findings were consistent across these approaches.

Despite the potential limitations of the CGT, we found some evidence, albeit weak, that lower risk taking might precede depressive symptoms in adolescent females. We interpreted this finding with caution, but it would be worth investigating in future studies with sufficient power to test interactions between depression and gender. Although we had a large sample, we might still have been under powered to detect an association in males. The effect size we observed in females was small and there were fewer males with exposure and outcome. It is also possible that low reward seeking is a risk factor for depressive symptoms in females but not in males.

Our finding of a large gender difference in risk adjustment at age 14 is consistent with adult studies. The finding suggests that males increase their bets more as the odds of winning increase. Our data suggests that this gender difference is not present at age 11. Interestingly, risk adjustment increased in males and females between 11 and 14, but to a larger extent in males. Adult gender differences in risk taking and reward seeking might emerge between the ages 11 and 14. In contrast to adult studies of the CGT, we found that, in addition to increasing their bets more as odds increased, adolescent males bet a larger proportion of their points than females (on average across all ratios). One possibility is that prior adult studies lacked statistical power to detect this difference. An alternative possibility is that adolescent gender differences in risk taking decline with age.

Our finding of no evidence for an association between risk taking to obtain reward and emotional/depressive symptoms is inconsistent with other cohort studies (Forbes et al., 2007; Rawal et al., 2013). Our sample was larger and we adjusted for a wider range of confounders, so it is possible that these prior findings are due to selection bias or confounding.

Our lack of evidence for cross-sectional and longitudinal associations between risk taking to obtain reward and depressive symptoms might have occurred for several reasons. First, risk taking to obtain reward, at least how it is assessed in the CGT, might have no association with depressive symptoms in children and adolescents. This is an important possibility, and one we cannot rule out. If so, it has implications for how we think about the aetiology, treatment and prevention of depressive symptoms in children and adolescents.

Second, the CGT conflates reward seeking and avoidance of punishment. Reduced betting, even when odds of winning are high, might occur because participants are less motivated by reward or because they want to avoid punishment. There is strong evidence, albeit in adults, that positive information processing is reduced in depression whilst negative information processing is unaltered (Lewis et al., 2017). It is possible that, in our study, the association between depressive symptoms and reward seeking was masked by conflation of positive and negative processing.

It is also possible that more self-relevant or emotional information is more important for emotional and depressive symptoms. In adult experimental psychopharmacology studies, antidepressants have been found to modify recall of self-relevant personality descriptors and recognition of emotional facial expressions (Harmer, Duman, & Cowen, 2017). Adolescence is a time when peer influences become more important. More socially relevant information might be more pertinent in the development of adolescent depressive symptoms.

## Key points

- We used a large nationally representative UK cohort to study adolescent risk taking (assessed with the Cambridge Gambling Task) and depressive symptoms and investigate gender differences
- We found that adolescent females were less risk taking than males
- We found no strong evidence that risk taking, assessed with this task, was associated with depressive symptoms
- Reduced risk taking may not be a risk factor for adolescent depression and the Cambridge Gambling Task is not a good candidate for future neuroimaging studies

## Acknowledgement

We are extremely grateful to the families who took part in the study, and the researchers and study personnel for all of their hard work. The Millennium Cohort Study is core funded by the Economic and Social Research Council (ESRC) and a consortium of Government departments. The authors gratefully acknowledge access to the Millennium Cohort Study data, provided by the Institute of Education, Centre for Longitudinal Studies.

