## Supplement for "Risk taking to obtain reward: gender differences and associations with emotional and depressive symptoms in a nationally representative cohort of UK adolescents"

**Means**

Supplementary Table 1. Mean (SD) SDQ scores at age 11, according to risk taking and risk adjustment in quintiles. Complete case sample (n=10,396).

| Risk taking quintiles | Mean (SD) | | | Risk adjustment quintiles | Mean (SD) | | |
| --- | --- | --- | --- | --- | --- | --- | --- |
|  | SDQ  overall | SDQ  females | SDQ  males |  | SDQ  overall | SDQ  females | SDQ  males |
| 1 (5-38) | 1.85 (1.93)  (n=2017) | 1.87 (1.91)  (n=1433) | 1.77 (1.98)  (n=584) | 1 (-.64 - -.09) | 2.04 (2.06)  (n=1997) | 2.10 (2.06)  (n=1034) | 1.97 (2.07)  (n=963) |
| 2 (39-47) | 1.82 (1.98)  (n=2199) | 1.83 (1.97)  (n=1216) | 1.80 (2.01)  (n=983) | 2 (-.08- -.35) | 1.87 (2.01)  (n=2028) | 2.03 (2.07)  (n=1019) | 1.72 (1.93)  (n=1009) |
| 3 (48-55) | 1.76 (1.94)  (n=1947) | 1.90 (2.01)  (n=978) | 1.63 (1.86)  (n=969) | 3 (.36-.78) | 1.79 (1.89)  (n=2065) | 1.83 (1.90)  (n=1029) | 1.74 (1.87)  (n=1036) |
| 4 (56-63,) | 1.80 (1.95)  (n=2197) | 2.01 (2.02)  (n=918) | 1.64 (1.89)  (n=1279) | 4 (.79-1.39) | 1.72 (1.93)  (n=2132) | 1.76 (1.89)  (n=1037) | 1.69 (1.96)  (n=1095) |
| 5 (64-95) | 1.86 (1.98)  (n=2036) | 1.95 (1.97)  (n=614) | 1.82 (1.98)  (n=1422) | 5 (1.40-6.00) | 1.68 (1.89)  (n=2174) | 1.80 (1.91)  (n=1040) | 1.57 (1.86)  (n=1134) |

Supplementary Table 2. Mean (SD) MFQ scores at age 14, according to risk taking and risk adjustment in quintiles. Complete case sample (n=8628).

| Risk taking quintiles | Mean (SD) | | | Risk adjustment quintiles | Mean (SD) | | |
| --- | --- | --- | --- | --- | --- | --- | --- |
|  | MFQ overall | MFQ females | MFQ males |  | MFQ overall | MFQ females | MFQ males |
| 1 (5-38) | 5.92  (n=1674) | 6.93  (n=1120) | 3.86  (n=554) | 1 (-.31-.021) | 5.79 (5.86)  (n=1608) | 7.12 (6.42)  (n=941) | 3.92 (4.31)  (n=667) |
| 2 (39-47) | 5.87  (n=1672) | 7.26  (n=969) | 3.96  (n=703) | 2 (-.22-.67) | 5.66 (5.94)  (n=1647) | 7.05 (6.54)  (n=803) | 4.05 (4.66)  (n=764) |
| 3 (48-55) | 5.70  (n=1903) | 7.27  (n=986) | 4.00  (n=917) | 3 (.68-1.14) | 5.43 (5.80)  (n=1712) | 6.89 (6.52)  (n=843) | 4.01 (4.58)  (n=869) |
| 4 (56-63,) | 5.35  (n=1652) | 7.06  (n=732) | 4.00  (n=920) | 4 (1.15-1.77) | 5.72 (6.04)  (n=1804) | 7.52 (6.80)  (n=872) | 4.04 (4.65)  (n=932) |
| 5 (64-95) | 4.99  (n=1727) | 6.80  (n=547) | 4.15  (n=1180) | 5 (1.78-5.10) | 5.26 (5.69)  (n=1857) | 6.83 (6.54)  (n=815) | 4.04 (4.57)  (n=1042) |

Supplementary Table 3. Mean (SD) MFQ scores at age 14, according to risk taking quintiles, at age 11. Complete case sample (n=8418).

| Risk taking | Mean (SD) | | | Risk adjustment | Mean (SD) | | |
| --- | --- | --- | --- | --- | --- | --- | --- |
|  | MFQ overall | MFQ females | MFQ males |  | MFQ overall | MFQ females | MFQ males |
| 1 (5-37) | 6.37 (6.31)  (n=1652) | 7.18 (6.62)  (n=1192) | 4.27 (4.81)  (n=460) | 1 (-.64 - -.09) | 5.65 (5.81)  (n=1518) | 7.07 (6.51)  (n=812) | 4.01 (4.34)  (n=706) |
| 2 (38-49) | 5.87 (5.97)  (n=1815) | 7.19 (6.66)  (n=1072) | 4.15 (4.56)  (n=788) | 2 (-.08- -.35) | 5.55 (5.79)  (n=1625) | 7.00 (6.31)  (n=836) | 4.03 (4.73)  (n=789) |
| 3 (50-57) | 5.46 (5.85) (n=1591) | 6.97 (6.69)  (n=793) | 3.95 (4.40)  (n=798) | 3 (.36-.78) | 5.48 (6.01)  (n=1679) | 6.97 (6.86)  (n=857) | 3.93 (4.48)  (n=822) |
| 4 (58-67) | 5.32 (5.65)  (n=1784) | 7.07 (6.55)  (n=758) | 4.03 (4.46)  (n=1026) | 4 (.79-1.39) | 5.53 (5.83) (n=1754) | 6.74 (6.51)  (n=853) | 4.38 (4.84)  (n=905) |
| 5 (68-95) | 4.89 (5.51)  (n=1576) | 6.68 (6.54)  (n=487) | 4.09 (4.77)  (n=1089) | 5 (1.40-6.00) | 5.74 (5.97)  (n=1838) | 7.53 (6.76)  (n=899) | 4.03 (4.50)  (n=939) |

**Risk adjustment as exposure variable**

Supplementary Table 4. Cross-sectional associations between risk adjustment (continuous exposure) and emotional symptoms at age 11 (continuous outcome), complete case sample (n=10,396).

| Sample overall | SDQ change for a 1-point increase in risk adjustment | P value |
| --- | --- | --- |
| Model 1a. Univariable (n=10,396) | -.13 (-.19 to -.08) | <.0001 |
| Model 1b. Model 1a adjusted^a^ | -.04 (-.09 to .01) | .121 |
| Model 1c. Model 1b adjusted for gender | -.04 (-.09 to .01) | .124 |
| Sub-group analyses by gender | SDQ change for a 1-point increase in risk adjustment | P value |
| Model 2a. Univariable, females (n= 5159) | -.14 (-.21 to -.07) | <.0001 |
| Model 2b. Model 2a adjusted^b^ | -.05 (-.13 to .02) | .128 |
| Model 3a. Univariable, males (n=5237) | -.13 (-.18 to -.07) | <.0001 |
| Model 3b. Model 3a adjusted^b^ | -.03 (-.09 to .04) | .436 |

^a^Adjusted for confounders measured at or as close as possible to the time of the exposure: family income, maternal education, child age, child ethnicity, child IQ, main carer depressive symptoms.

^b^Adjusted for the above confounders and, in addition, stage of breast development in females or stage of facial hair development in males.

Supplementary Table 5. Cross-sectional associations between risk adjustment (continuous exposure) and depressive symptoms at age 14 (continuous outcome), complete case sample (n=8628).

| Sample overall | MFQ change for a 1-point increase in risk adjustment | P value |
| --- | --- | --- |
| Model 1a. Univariable (n=8628) | -.17 (-.33 to -.02) | .029 |
| Model 1b. Model 1a adjusted^a^ | -.10 (-.26 to .06) | .205 |
| Model 1c. Model 1a adjusted for sex | .03 (-.13 to .19) | .699 |
| Sub-group analyses by gender | MFQ change for a 1-point increase in risk adjustment | P value |
| Model 2a. Univariable, females (n=4354) | -.07 (-.33 to .19) | .588 |
| Model 2b. Model 2a adjusted^b^ | .05 (-.22 to .32) | .714 |
| Model 3a. Univariable males (n=4274) | -.01 (-.18 to .15) | .883 |
| Model 3b. Model 3a adjusted^b^ | .01 (-.16 to .18) | .905 |

^a^Adjusted for confounders measured at or as close as possible to the time of the exposure: family income, maternal education, child age, child ethnicity, child IQ, main carer depressive symptoms.

^b^ Adjusted for the above confounders and, in addition, stage of breast development in females or stage of facial hair development in males.

Supplementary Table 6. Longitudinal association between risk adjustment (continuous exposure) at age 11 and depressive symptoms (continuous outcome) at age 14, complete cases (n=8418).

| Sample overall | MFQ change for a 1-point increase in risk adjustment | P value |
| --- | --- | --- |
| Model 1a. Univariable (n= 8,418) | -.03 (-.18 to .12) | .666 |
| Model 1b. Model 1a adjusted^a^ | .09 (-.07 to .25) | .275 |
| Model 1c. Model 1b adjusted for sex | .12 (-.03 to .28) | .123 |
| Sub-group analyses by gender | MFQ change for a 1-point increase in risk adjustment | P value |
| Model 2a. Univariable, females (n=4257) | -.03 (-.30 to .24) | .810 |
| Model 2b. Model 2a adjusted^b^ | .07 (-.22 to .35) | .635 |
| Model 3a. Univariable, males (n=4161) | .01 (-.15 to .18) | .858 |
| Model 3b. Model 3a adjusted^b^ | .15 (-.01 to .32) | .069 |

^a^Adjusted for confounders measured at or as close as possible to the time of the exposure: family income, maternal education, main carer depressive symptoms, child age, child ethnicity, child IQ and SDQ total difficulties score.

^c^Adjusted for the above confounders and, in addition, stage of breast development in females or stage of facial hair development in males.

**Imputed associations with risk taking**

Supplementary Table 7. Cross-sectional associations between risk taking (continuous exposure) and emotional symptoms at age 11 (continuous outcome), multiply imputed sample (n=12,355).

| Sample overall | SDQ change for a 20-point increase in risk taking | P value |
| --- | --- | --- |
| Model 1 Univariable (n=12,355) | -.01 (-.07 to .04) | .617 |
| Model 2: Model 1 adjusted^a^ | -.03 (-.08 to .02) | .221 |
| Model 3: Model 2 adjusted for sex | -.03 (-.08 to .03) | .320 |
| Sub-group analyses by gender | SDQ change for a 20-point increase in risk taking | P value |
| Model 4: Univariable, females (n=6144) | .02 (-.05 to .10) | .522 |
| Model 5: Model 4 adjusted^b^ | -.03 (-.09 to .04) | .445 |
| Model 6: Univariable males (n=6211) | .02 (-.06 to .09) | .646 |
| Model 7: Model 6 adjusted^b^ | -.03 (-.10 to .05) | .481 |

^a^Adjusted for confounders measured at or as close as possible to the time of the exposure: family income, maternal education, child age, child ethnicity, child IQ, main carer depressive symptoms.

^b^Adjusted for the above confounders and, in addition, stage of breast development in females and stage of facial hair development in males.

Supplementary Table 8. Cross-sectional associations between risk taking (continuous exposure) and depressive symptoms at age 14 (continuous outcome), imputed sample (n=10,578).

| Sample overall | MFQ change for a 20-point increase in risk taking | P value |
| --- | --- | --- |
| Model 1 Univariable (n=10,578) | -.58 (-.75 to -.41) | <.0001 |
| Model 2: Model 1 adjusted^a^ | -.44 (-.62 to -.26) | <.0001 |
| Model 3: Model 2 adjusted for sex | -.01 (-.19 to .17) | .928 |
| Sub-group analyses by gender | MFQ change for a 20-point increase in risk taking | P value |
| Model 4: Univariable, females (n=5324) | -.05 (-.38 to .28) | .776 |
| Model 5: Model 4 adjusted^b^ | -.08 (-.41 to .24) | .605 |
| Model 6: Univariable males (n=5254) | .05 (-.17 to .26) | .649 |
| Model 7: Model 6 adjusted^b^ | .05 (-.17 to .26) | .679 |

^a^Adjusted for confounders measured at or as close as possible to the time of the exposure: family income, maternal education, child age, child ethnicity, child IQ, main carer depressive symptoms.

^b^ Adjusted for the above confounders and, in addition, stage of breast development in females and stage of facial hair development in males.

Supplementary Table 9. Longitudinal association between risk taking (continuous exposure variable) at age 11 and depressive symptoms (continuous outcome) at age 14, imputed sample (n=12,355).

| Sample overall | MFQ change for a 20-point increase in risk taking | P value |
| --- | --- | --- |
| Model 1 Univariable (n=12,355). | -.60 (-.76 to -.43) | <.0001 |
| Model 2: Model 1 adjusted^a^ | -.30 (-.46 to -.14) | <.0001 |
| Model 3: Model 2 adjusted for sex | -.13 (-.29 to .03) | .112 |
| Sub-group analyses by gender | MFQ change for a 20-point increase in risk taking | P value |
| Model 4: Univariable, females (n=6144) | -.21 (-.44 to .03) | .089 |
| Model 5: Model 4 adjusted^b^ | -.24 (-.47 to -.00) | .047 |
| Model 6: Univariable males (n=6211) | .06 (-.15 to .26) | .583 |
| Model 7: Model 6 adjusted^b^ | -.01 (-.20 to .19) | .936 |

^a^Adjusted for confounders measured at or as close as possible to the time of the exposure: family income, maternal education, main carer depressive symptoms, child age, child ethnicity, child IQ and SDQ total difficulties score.

^c^Adjusted for the above confounders and, in addition, stage of breast development in females and stage of facial hair development in males.

**SDQ at 14 years of age as outcome**

Supplementary Table 10. Cross-sectional associations between risk taking (continuous exposure) and emotional symptoms at age 14 (continuous outcome), complete case sample.

| Sample overall | SDQ change for a 20-point increase in risk taking | P value |
| --- | --- | --- |
| Model 1a. Univariable (n=8431) | -.14 (-.21 to -.07) | <.0001 |
| Model 1b. Model 1a adjusted^a^ | -.17 (-.23 to -.10) | <.0001 |
| Model 1c. Model 1b adjusted for gender | -.06 (-.12 to .01) | .091 |
| Sub-group analyses by gender | SDQ change for a 20-point increase in risk taking | P value |
| Model 2a. Univariable, females (n=4238) | -.08 (-.19 to .03) | .159 |
| Model 2b. Model 2a adjusted^b^ | -.12 (-.23 to -.02) | .022 |
| Model 3a. Univariable males (n=4193) | .04 (-.06 to .13) | .428 |
| Model 3b. Model 3a adjusted^b^ | -.00 (-.09 to .09) | .981 |

^a^Adjusted for confounders measured at or as close as possible to the time of the exposure: family income, maternal education, child age, child ethnicity, child IQ, main carer depressive symptoms.

^b^ Adjusted for the above confounders and, in addition, stage of breast development in females and stage of facial hair development in males.

Supplementary Table 11. Longitudinal association between risk taking (continuous exposure variable) at age 11 and emotional symptoms (continuous outcome) at age 14, complete cases.

| Sample overall | MFQ change score for a 20-point increase in risk taking | P value |
| --- | --- | --- |
| Model 1 Univariable (n= 8,205). | -.04 (-.11 to .03) | .305 |
| Model 2: Model 1 adjusted^a^ | -.17 (-.23 to -.11) | <.0001 |
| Model 3: Model 2 adjusted for sex | -.04 (-.10 to .03) | .254 |
| Sub-group analyses by gender | MFQ change for a 20-point increase in risk taking | P value |
| Model 4: Univariable, females (n=4139) | .04 (-.06 to .14) | .454 |
| Model 5: Adjusted females^b^ | -.06 (-.15 to .03) | .174 |
| Model 6: Univariable males (n= 4,066) | .12 (.02 to .21) | .015 |
| Model 7: Adjusted males^b^ | -.01 (-.09 to .07) | .841 |

^a^Adjusted for confounders measured at or as close as possible to the time of the exposure: family income, maternal education, main carer depressive symptoms, child age, child ethnicity, child IQ and SDQ total difficulties score.

^c^Adjusted for the above confounders and, in addition, stage of breast development in females and facial hair development in males.
